# Weak representation of awake/sleep states by local field potentials in aged mice

**DOI:** 10.1101/2021.10.13.464191

**Authors:** Daichi Konno, Yuji Ikegaya, Takuya Sasaki

## Abstract

Senescence affects various aspects of sleep, and it remains unclear how sleep-related neuronal network activity is altered by senescence. Here, we recorded local field potential signals from multiple brain regions covering the forebrain in young (10-week-old) and aged (2-year-old) mice. Interregional LFP correlations across these brain regions showed smaller differences between awake and sleep states in aged mice. Multivariate analyses with machine learning algorithms with uniform manifold approximation and projection (UMAP) and robust continuous clustering (RCC) demonstrated that these LFP correlational patterns in aged mice less represented awake/sleep states than those in young mice. By housing aged mice in an enriched environment, the LFP patterns were restored to those observed in young mice. Our results demonstrate senescence-induced changes in neuronal activity at the network level and provide insight into the prevention of pathological symptoms associated with sleep disturbance in senescence.

## Introduction

Senescence reduces the quality of sleep (e.g., shorter sleep time, longer sleep-onset latency, sleep fragmentation, and insomnia) ^1,2,3^. Sleep disturbance in senescence increases the onset risk of many diseases, such as Alzheimer’s disease, depression, and diabetes ^4^. Many biological factors at the molecular and cellular levels in the aged brain have been revealed, such as the loss of neurotransmitters ^5^, synapses ^6,7^, and neurons ^8^. The integration of these microscopic mechanisms is considered to reduce brain volume ^9,10,11,12^ and disrupt neuronal network activity that sustains normal awake/sleep cycles.

Physiological studies have demonstrated that cortical brain oscillations in specific frequency bands are reduced by aging ^13,14,15,16^. These observations suggest that neuronal network mechanisms to create sleep-related oscillatory signals are degraded in senescence. To further understand sleep-related brain network activity in senescence,, functional connectivity (i.e., correlated activity) patterns across multiple brain regions need to be considered. As sleep states are maintained through interactions with many, not single, brain regions in various frequency bands ^17^, investigations from multiple brain areas and multiple frequency bands are necessary to comprehend senescence-related neuronal network activity. In addition, investigations with temporal windows associated with dynamic changes between awake/sleep states, rather than entire data sampling periods, are necessary to precisely capture and evaluate senescence-related neuronal network activity.

To address these issues, this study compared awake/sleep-related neuronal network activity between young (10 week) and aged (2 year) mice by applying correlational and multivariate analyses to local field potential (LFP) data recorded from multiple brain regions covering the entire forebrain area. After identifying decreases in dynamic changes in LFP signals that differentiate awake/sleep states in aged mice, we assessed whether such awake/sleep-related brain activity can be restored by housing them in an enriched environment (EE).

## Results

### Sleep time decreases in aged mice

We established a recording system that simultaneously monitors LFP signals from six brain regions and dorsal neck EMG signals in freely moving mice using a custom-made plastic device (Fig. 1A). The six brain regions included the anterior cingulate cortex (ACC), primary motor cortex (M1), striatum (STR), primary somatosensory cortex (S1), hippocampus (HPC), and medial parietal association cortex (MtPA) (Fig. 1B). These brain areas were selected so that they covered the wide range of the forebrain from anterior to posterior parts. Recording sites were verified by histological inspection. Figure 1C shows representative simultaneous recordings of LFP traces and an EMG trace obtained from a freely moving mouse. These electrophysiological signals were recorded from young (8- to 10-week-old; collectively termed 10-week) and aged (95- to 105-week-old; collectively termed 2-year) mouse groups.

**Figure 1.**
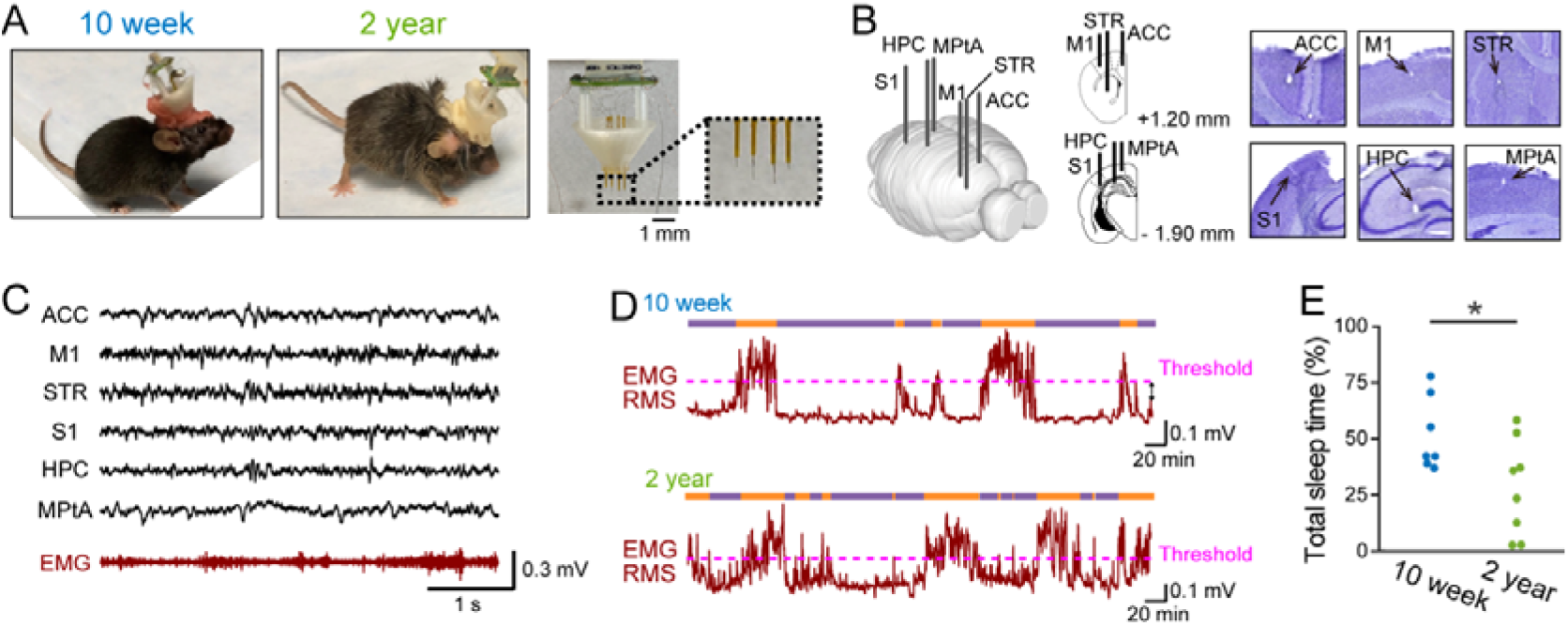
Multisite LFP recordings and definition of sleep states in aged mice. **(A)** (Left) A 10-week and 2-year mouse implanted with an electrode array. (Right) A custom-made electrode array. The dotted box is magnified in the right inset, showing electrodes protruding from the electrode assembly with a length of 0.5–2.3 mm to be inserted into the brain. **(B)** (Left) Electrodes implanted into six brain regions. (Right) Histological confirmation of an electrode in each region. **(C)** Representative LFP signals recorded from the six brain regions (black) and an EMG signal (brown). **(D)** The RMS (magnitude) of EMG signals in 10-week (top) and 2-year (bottom) mice. The magenta dotted lines represent the detection thresholds of 1.5 SD above the mean. The upper bars represent sleep (purple) and awake (orange) states. **(E)** The percentage of sleep periods defined from EMG signals to total recording periods (*n* = 7 and 8 mice). Each dot represents an individual mouse. \**P* < 0.05, Student’s *t*-test.

First, we examined how EMG-based awake/sleep states differed between these mouse groups. EMG signals were converted to root mean square (RMS) traces with a bin size of 1 s (Fig. 1D). Sleep states were defined as periods when EMG signals were below a threshold of the mean value of the EMG signal plus 1.5 SD of the EMG signal for more than 180 s. In the representative 10-week and 2-year mice shown in Figure 1D, sleep time accounted for 67.3% and 50.2% of the total recording time, respectively. Overall, the percentage of sleep time in 2-year mice was significantly lower than that in 10-week mice (Fig. 1E; *n*□=□7 and 8 mice; *t*_13_ = 2.39, *P* = 0.033, Student’s *t*-test).

### Smaller differences in interregional LFP correlations between awake and sleep states in aged mice

We next analyzed LFP signals from multiple brain regions. For each brain region, an LFP signal was converted to its power traces in six frequency bands (delta (δ), 1–4 Hz; theta (θ), 4–8 Hz; alpha (α), 8–13 Hz; beta (β), 13–30 Hz; low-gamma (low-γ (L)), 30–50 Hz; high-gamma (high-γ (H)), 50–100 Hz) (Fig. 2A). First, we compared LFP power in these individual frequency bands averaged over all awake and sleep states. We found no significant differences between 10-week and 2-year mice in average LFP power in most of the frequency bands and brain regions (Supplementary Fig. 1).

**Figure 2.**
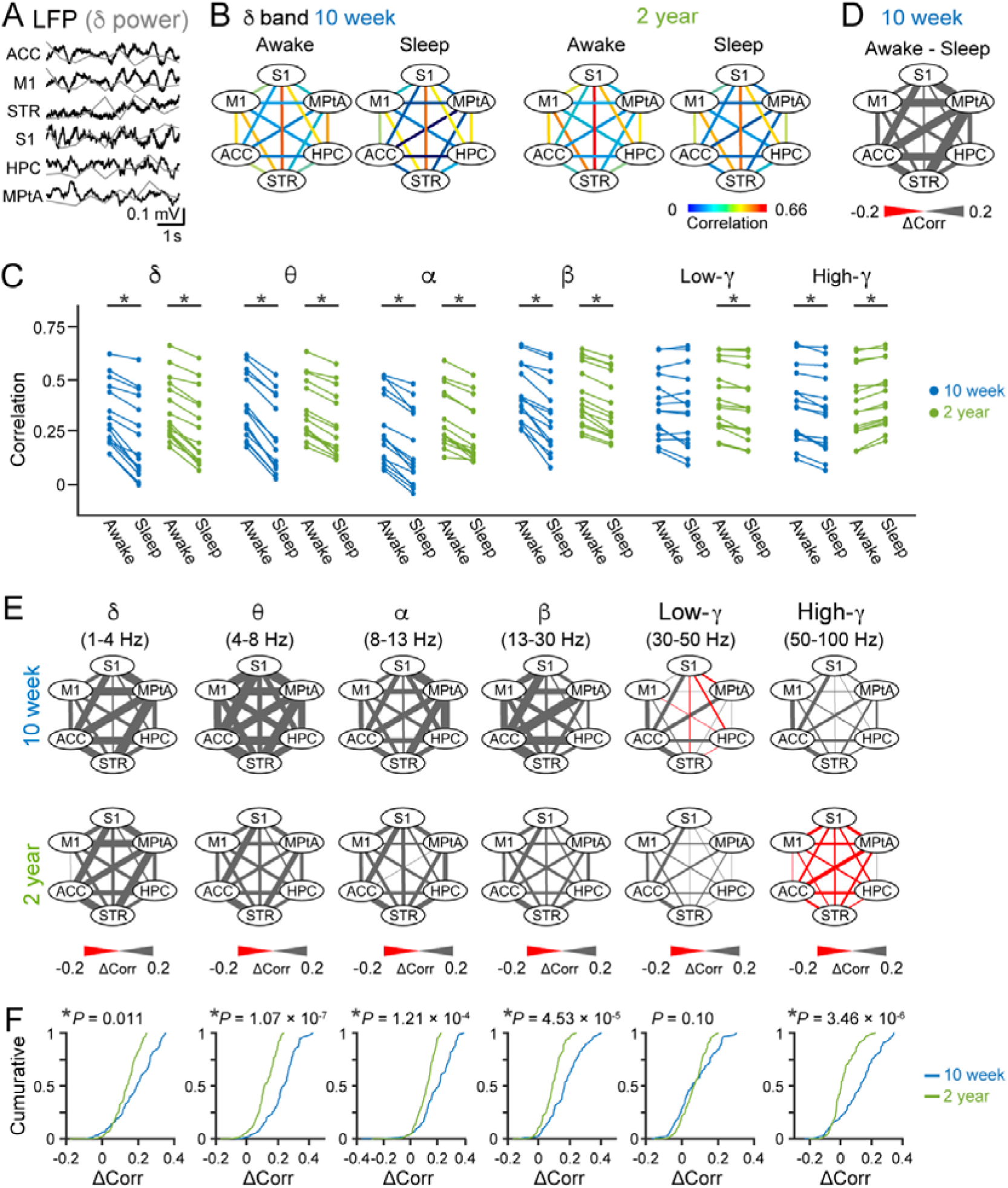
Interregional correlations of LFP power changes across awake/sleep states in young and aged mice. **(A)** LFP signals were converted into normalized delta power (superimposed as gray lines) every 1 s. **(B)** Representative color-coded maps showing correlation coefficients of delta power changes for 15 pairs of the six brain regions. Data are from a representative mouse at each age. **(C)** Comparisons of averaged correlational power changes in six frequency bands between awake and sleep states. Each line represents each brain region pair. \**P* < 0.05, paired *t*-test. **(D)** A color-coded map showing differences in the power correlations (Δcorr) between awake and sleep states, constructed from B. **(E)** Color-coded maps showing Δcorr in awake and sleep states in six frequency bands. Data are averaged from all mice. **(F)** Cumulative distributions of Δcorr. The *P* value between 10-week and 2-year mice defined by the Mann–Whitney U test is described above.

We next examined regional correlations of LFP power changes (i.e., functional connection) in awake/sleep states. To obtain an overview of functional connections from all datasets, we computed correlation coefficients of LFP power changes in individual frequency bands between two brain regions throughout the entire recording session. Correlation coefficients from all brain region pairs (_6_C_2_ = 15) from a representative mouse are summarized as a color-coded map and shown in Figure 2B. In both the 10-week and 2-year mice, the correlation coefficients in all the pairs during awake states were significantly higher than those during sleep states in the majority of frequency bands (Fig. 2C; *n* = 7 and 8 mice; *P* = 1.41 × 10^-6^ (10-week, delta), *P* = 7.23 × 10^-11^ (10-week, theta), *P* = 2.25 × 10^-8^ (10-week, alpha), *P* = 1.66 × 10^-6^ (10-week, beta), *P* = 0.067 (10-week, low-gamma), *P* = 3.38 × 10^-5^ (10-week, high-gamma), *P* = 6.62 × 10^-8^ (2-year, delta), *P* = 1.90 × 10^-8^ (2-year, theta), *P* = 6.19 × 10^-8^ (2-year, alpha), *P* = 7.33 × 10^-9^ (2-year, beta), *P* = 1.76 × 10^-4^ (2-year, low-gamma), *P* = 2.77 × 10^-6^ (2-year, high-gamma), paired *t*-test). While significant differences between awake and sleep states were found in both of the mouse groups, the differences in the 10-week group appeared larger than those in the 2-year group (Fig. 2C, as indicated by the blue lines). To test this idea, differences in correlation coefficients between awake and sleep states were calculated as Δcorr in individual brain region pairs and summarized as a color-coded map (Fig. 2D). Figure 2E shows all maps of Δcorr in the six frequency bands averaged over all mice. Overall, cumulative distributions confirmed significantly larger Δcorr in the 10-week group than in the 2-year group in all frequency bands, except the low-gamma band (Fig. 2F; *n* = 7 and 8 mice; *P* = 0.011 (delta), *P* = 1.07 × 10^-7^ (theta), *P* = 1.21 × 10^-4^ (alpha), *P* = 4.53 × 10^-5^ (beta), *P* = 0.10 (low-gamma), *P* = 3.46 × 10^-6^ (high-gamma), Kolmogorov–Smirnov test). These results demonstrate that regional correlations of LFP power changes more strongly differentiate awake and sleep states in young mice than in aged mice.

### Weak representation of awake/sleep states by multi-region LFP correlations in aged mice

The analyses in Figure 2 focused on overall differences in LFP power averaged over all awake/sleep periods. Under natural conditions, awake/sleep states continuously vary across time (generally tens of seconds to several minutes), and a crucial issue is whether brain LFP patterns at specific time periods could differentiate such time-varying awake/sleep state patterns. Recording periods of 2.5–3.5 hours were divided into bins of 10 s, and each bin was categorized as an awake or a sleep bin, depending on whether the bin contained an EMG RMS exceeding the threshold for more than 1 s. In each bin, correlation coefficients of LFP power changes in each frequency band were computed across the 15 pairs of the six brain regions to obtain a 15-dimensional vector (Fig. 3A). Our analysis tested whether these correlational LFP patterns in each bin could represent an awake or sleep bin. For each frequency band, we applied UMAP (uniform manifold approximation and projection), a dimension reduction technique without any subjective bias, to these vectors. Figure 3B shows representative two-dimensional UMAP plots in the delta band for a 10-week and 2-year mouse. In these graphs, the plots appeared to be somewhat separated between awake and sleep bins. To quantify the degree of separation, the robust continuous clustering (RCC) algorithm was applied to all the plots in each graph. After identifying clusters by the RCC, each cluster was categorized as an awake or a sleep cluster, depending on whether there were more awake or sleep bins in the cluster, respectively. An F1 score was then computed as an index based on the percentage of awake/sleep bins assigned in awake/sleep clusters (Fig. 3B and 3C). To assess the significance of an F1 score, shuffled datasets were constructed by randomly shuffling bin types (awake/sleep) to which the individual plots across all the plots were categorized. An F1 score was considered to be significant when it was higher than all F1 scores computed from 1000 shuffled datasets (a distribution shown in the bottom panel in Figure 3B). In the 10-week mice, 71.4% of F1 scores were significant in all single frequency bands (Fig. 3C, top; indicated by the closed circles), suggesting that regional correlations of LFP power changes computed in a single frequency band, to some extent, represent awake/sleep states. On the other hand, the percentages decreased to 12.5–50.0% in the 2-year mice (Fig. 3C, bottom; indicated by the closed circles), suggesting weaker representations of awake/sleep states by LFP correlations in aged mice. Next, we asked whether the awake/sleep separations in LFP patterns were improved when the datasets in all six frequency bands were employed in an analysis. All vectors in the six frequency bands in individual bins were combined to construct 90-dimensional vectors (Fig. 3D), and the same procedures, plotting on UMAP dimensions and computing F1 scores, were applied to these 90-dimensional vectors (Fig. 3E). All 10-week mice showed significant F1 scores (Fig. 3C, top; indicated by the closed circles), whereas only 20% of the 2-year mice showed significant F1 scores (Fig. 3C, bottom; indicated by the closed circles). Overall, F1 scores were significantly higher in the 10-week mice than in the 2-year mice (Fig. 3F; *n*□=□7 and 8 mice; *t_13_* = 5.00, *P* = 2.43 × 10^-4^, Student’s *t*-test). Taken together, our multidimensional analyses demonstrate that awake and sleep states are less precisely represented by LFP correlational patterns in 2-year mice than in 10-week mice.

**Figure 3.**
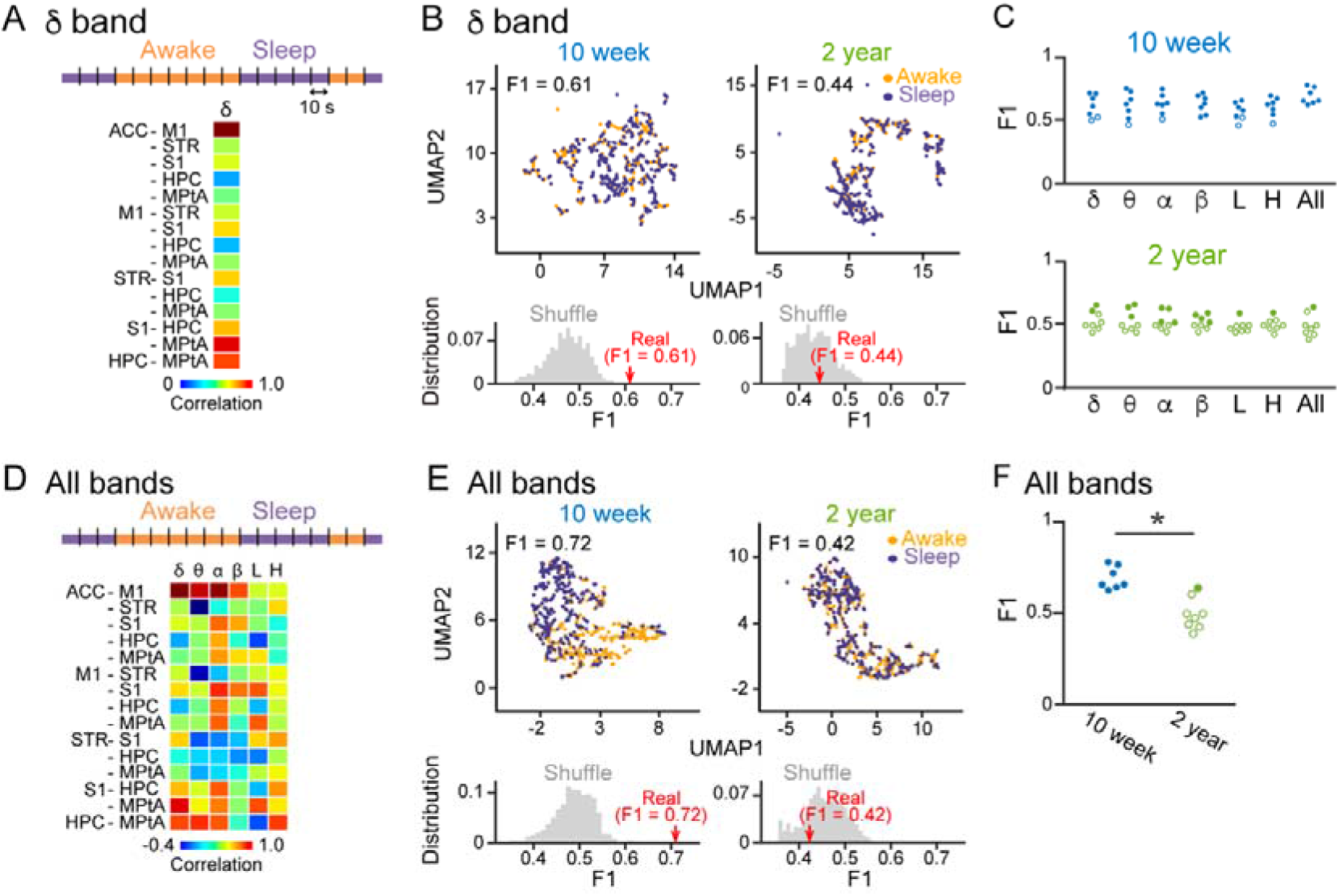
LFP correlational patterns less represented awake and sleep states in aged mice. **(A)** Schematic of analysis. Each 10-s bin was categorized as an awake (orange) or a sleep (purple) bin based on EMG signals, and correlations of power changes in a frequency band across the 15 region pairs were concatenated in a column. **(B)** (Top) Visualization of single-mouse data by UMAP plots. Orange and purple dots represent awake and sleep 10-s bins, respectively. F1 scores to quantify the separation of awake/sleep states from LFP patterns are shown above. (Bottom) The F1 score shown in the top panel (real, red arrow) was compared with a distribution of F1 scores computed from 1000 shuffled datasets in which awake/sleep states of all the plots were randomly shuffled (gray distribution). **(C)** F1 scores computed in individual frequency bands (left six bands) and all six bands combined (rightmost). Each dot represents an individual mouse. The closed and open circles represent significant and nonsignificant data, respectively. **(D,E)** Same as A and B, but correlations in the six frequency bands were used for the UMAP analysis. **(F)** Comparison of F1 scores in all six frequency bands between young and aged mice. Data are similar to those shown in C. \**P* < 0.05, Student’s *t*-test.

### Enriched environment restores LFP correlations to represent awake/sleep states in aged mice

For rodent animals, housing in an enriched environment (EE) has been shown to exert positive effects on various behavioral patterns and sleep quality ^18,19^. Here, we examined whether living in an EE could restore aging-related LFP changes (Fig. 4A). First, we tested whether our experimental EE conditions were sufficient to affect memory performance. An object location test was utilized in which a mouse was first introduced to two identical objects in an open field (acquisition phase) and then exposed to the same two objects, one of which was displaced to a new location (test phase) (Fig. 4B). If the mouse remembered the locations of the objects in the acquisition phase, it would spend more time exploring the object in the new position. The location index, representing the interaction time with the novel object in the test phase, was significantly higher than the chance level (50%) in 10-week mice (*n* = 6 mice; *t*_5_ = 2.48, *P* = 0.028, Student’s *t*-test versus 0.5) but not in 2-year mice without EE (termed 2-year non-EE mice; *n* = 9 mice; *t*_8_ = 0.19, *P* = 0.43, Student’s *t*-test versus 0.5) (Fig. 4B). On the other hand, 2-year mice housed in EE (termed 2-year EE mice) for more than 2 weeks exhibited a location index that was significantly higher than chance (*n* = 9 mice; *t*_8_ = 2.26, *P* = 0.027, Student’s *t*-test versus 0.5). These results confirmed that our EE conditions could restore spatial memory performance, which was decreased in aged mice. Furthermore, we confirmed gene expression patterns in the dorsal hippocampus in these mouse groups. Aged mice exhibited altered expression patterns of a subset of genes related to neuron development and synaptic signaling (Supplementary Figs. 2A, 2C and 3). On the other hand, the expression patterns in the 2-year EE mice were more similar to those observed in 10-week mice (Supplementary Fig. 2B), confirming the restoration of aging-induced gene expression by EE. EE tended to, although the difference was not significant, alter total sleep time, which was decreased in aged mice (Fig. 4C; *n*□=□7, 8, and 6 mice; *P* > 0.05, Tukey’s test).

**Figure 4.**
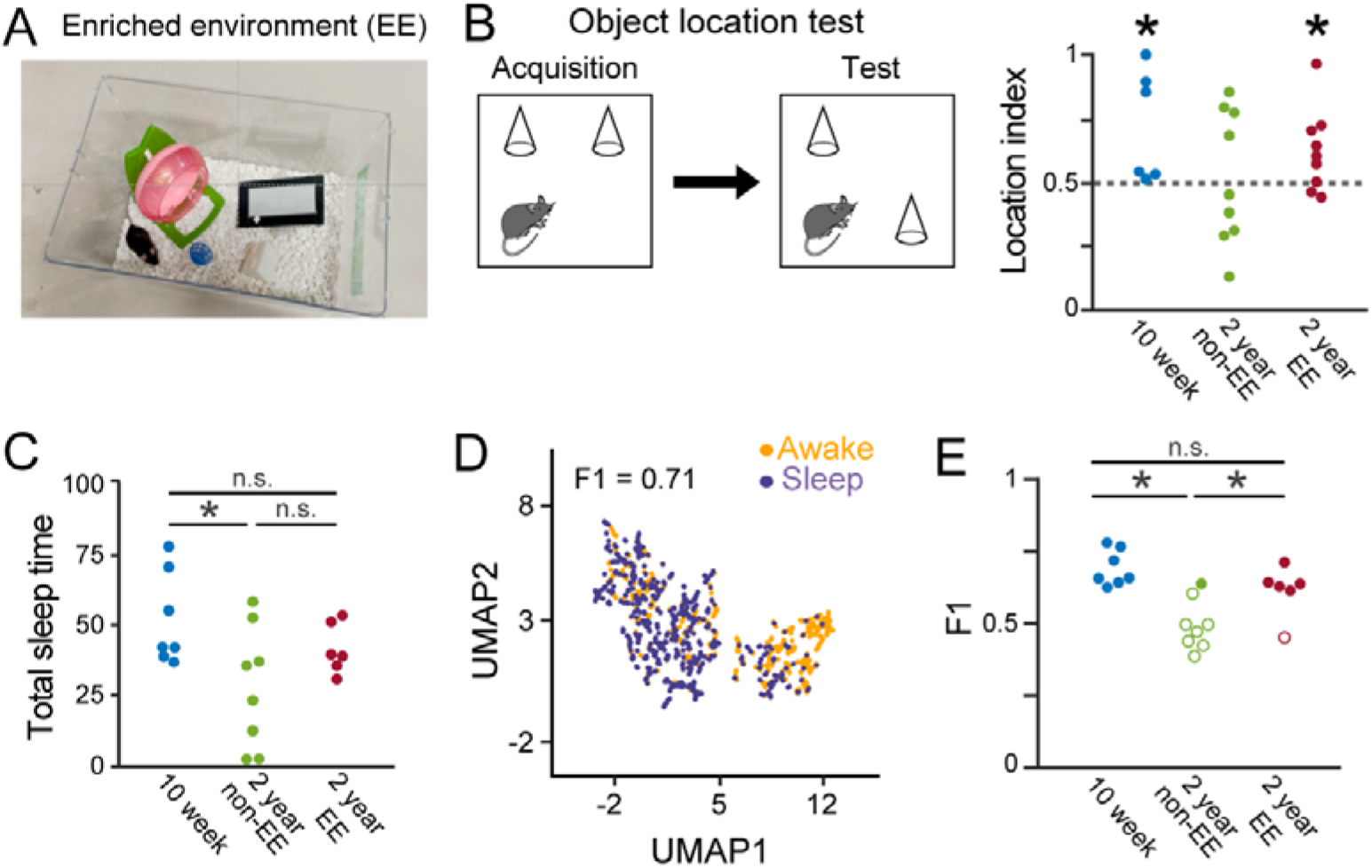
EE restores sleep-related neuronal activity in aged mice. **(A)** A photograph of the EE. **(B)** (Left) Schematic diagram of an object location test. (Right) Location index in the object location test (*n* = 6, 9, and 9 mice). Each dot represents an individual mouse. \**P* < 0.05, Student’s *t*-test versus 0.5 (dotted line). **(C)** The percentage of sleep periods to total recording periods (*n* = 7, 8, and 6 mice). The plots in the 10-week and 2-year non-EE groups are similar to those shown in Figure 1E, presented for comparison. \**P* < 0.05, Tukey’s test. **(D)** Same as Figure 3E but for a representative 2-year EE mouse. **(E)** Comparisons of F1 scores across the three mouse groups (*n* = 7, 8, and 5 mice). Each dot represents an individual mouse. The closed and open circles represent significant and nonsignificant data, respectively. The plots in the 10-week and 2-year non-EE groups are similar to those shown in Figure 3F, presented for comparison. \**P* < 0.05, Tukey’s test.

Next, we applied the same analytical procedures as in Figure 3F, plotting on UMAP dimensions and computing F1 scores, to the LFP datasets in all the frequency bands from the 2-year EE mice (Fig. 4D). Out of 6 mice tested, 5 (83.3%) mice exhibited significant F1 scores in the 2-year EE group, which was prominently higher than that (12%) in the 2-year non-EE group (Fig. 4E). Overall, the F1 scores in the 2-year EE group were significantly higher than those in the 2-year non-EE group (*n*□=□8 and 6; *P* < 0.05, Tukey’s test) and not significantly different from those in the 10-week group (*n*□=□7 and 6 mice; Tukey’s test). These results suggest that housing aged mice in EE conditions restores brain network LFP patterns to more precisely represent awake/sleep states.

## Discussion

To understand senescence-induced changes in neuronal activity at the network level, we developed a device that simultaneously records local field potential signals from multiple brain regions in the forebrain together with EMG signals by improving our previous methods ^20,21,22^. Consistent with a previous study ^23^, we confirmed that the total sleep time was shorter in aged mice. While some studies have shown that sleep time becomes longer in aged mice ^24,25^, this discrepancy may be due to experimental conditions such as novelty of recording environments for mice, recording time, or definition of sleep ^23,24,25^. We demonstrated that awake/sleep differences in interregional LFP correlations in the delta to high-gamma bands in the forebrain are smaller in aged mice. Machine learning algorithms with UMAP and RCC revealed that these LFP correlational patterns less represented awake/sleep states in aged mice.

In both young and aged mice, LFP power in single frequency bands in single brain regions was not sufficient to differentiate awake/sleep states but regional correlations of LFP power changes (i.e., functional connections) showed differences in awake/sleep states. The importance of focusing on interregional activity patterns is consistent with the suggestions that awake and sleep brain states are not simply sustained by activity levels of single brain regions but generated from complex interactions across widespread brain regions ^26,27^. When the datasets in all six frequency bands were combined, the representations of awake/sleep states by LFP signals were improved in young mice. In addition, both young and aged mice exhibited reduced correlations of LFP power changes across brain regions during sleep states compared with awake states, indicating decreased interregional information transfer and integration during sleep states. These implications are likely explained by the observation that functional connection patterns, defined based on correlated activity changes, decrease and become more independent modules in sleep states ^28,29^.

Our results demonstrated that interregional correlational patterns of LFP signals differentiated awake and sleep states less in aged mice than in young mice. Several molecular and cellular factors have been reported to be involved in aging, including decreases in neuromodulators ^5^, reduced expression of neuromodulator receptors ^30^, and decreased synchronicity of neurons in the suprachiasmatic nucleus ^31^. These mechanisms may cooperatively obscure the neuronal activity that cannot clearly distinguish awake and sleep states.

Finally, our results demonstrated that housing aged mice in EE conditions reinstated the aging-related decreases in LFP patterns such that they more precisely represented awake/sleep states. In addition to the positive effects of EE on various behavioral patterns ^18,19^ and gene expression patterns ^32,33^ reported previously, our study verified that EE can restore awake/sleep-related neuronal activity patterns. Considering that declines in sleep quality in senescence are crucial risk factors for neurodegenerative and psychiatric diseases ^4^, EE is expected to prevent these pathological symptoms through improvements in the quality of sleep and sleep-related neuronal activity.

## Materials and methods

### Ethical approvals

Animal experiments were performed with the approval of the Animal Experiment Ethics Committee at The University of Tokyo (approval number: P29-14) and according to the University of Tokyo guidelines for the care and use of laboratory animals. These experimental protocols were carried out in accordance with the Fundamental Guidelines for Proper Conduct of Animal Experiment and Related Activities in Academic Research Institutions (Ministry of Education, Culture, Sports, Science and Technology, Notice No. 71 of 2006), the Standards for Breeding and Housing of and Pain Alleviation for Experimental Animals (Ministry of the Environment, Notice No. 88 of 2006) and the Guidelines on the Method of Animal Disposal (Prime Minister’s Office, Notice No. 40 of 1995). All efforts were made to minimize the animals’ suffering.

### Animals

Male 8-to-10-week-old C57BL/6 mice (SLC, Shizuoka, Japan) and male 95-to-105-week-old C57BL/6 mice (Charles River Japan, Kanagawa, Japan) with a preoperative weight of 30 g were housed under conditions of controlled temperature and humidity (22 ± 1 °C, 55 ± 5%), maintained on a 12:12-h light/dark cycle (lights off from 07:00 to 19:00) with *ad libitum* access to food and water. All mice were housed individually.

### Enriched environment

All young 2-week-old mice were housed in a homecage. Two-year-old mice were randomly selected and exposed to either standard environment (non-EE) or enriched environment (EE) conditions for more than two weeks. For non-EE conditions, standard laboratory cages were used (24 cm × 17 cm × 12 cm), whereas mice maintained under EE conditions were housed in a rat cage (42 cm × 26 cm × 20 cm). The EE cages were equipped with a running wheel, a seesaw, a ball, and a climbing platform (Fig. 4A).

### Object location test

The experimental apparatus used in this study was an open-field box (47 cm × 37 cm with walls with a height of 31 cm). The apparatus was placed in a sound-isolated room. Identical plastic cones (10 cm in height × 6 cm in diameter) created using a 3D printer (Form 2, Formlabs, MA) were used as objects. This test consisted of a habituation day and a test day. On a habituation day, the mice were allowed to freely explore the apparatus without objects for 1 hour. On the next day, which was the test day, the same habituation procedure was first conducted for 30 min before starting the acquisition phase. In the acquisition phase, the mice were allowed to freely explore two identical objects that were placed symmetrically in the experimental apparatus for 5 min. The mouse was then removed from the apparatus and returned to its home cage. The objects were thoroughly cleaned with 70% ethanol. The open field box was cleaned with dry paper after each trial to ensure that it was saturated with the smell of the animals. A test phase was conducted 1 h after the acquisition phase. In the test phase, one of the objects (A2) was moved to a different location (A2′(D)), and the other object (A1) remained in the same position (A1(ND)) as in the acquisition phase. The terms D and ND indicate the displaced and the nondisplaced objects in the test phase, respectively. In the test phase, the mouse was allowed to freely explore the experimental apparatus for 5 min. Their behavior was recorded with a video camera mounted above the apparatus. The time spent exploring each object was measured using DeepLabCut ^34^. Exploration of an object was defined as pointing the nose toward the object at a distance of < 1 cm and/or touching it with the nose. To analyze memory performance, a location index was calculated as follows: TA2′(D)/(TA1(ND)+TA2′(D)), where TA2′(D) is the time exploring the displaced object and TA1(ND) is the time exploring the nondisplaced object.

### Preparation of an electrode array

An electrode array for brain LFP recording was assembled consisting of custom-made parts (Fig. 1A) and an electrical interface board (EIB) (Neuralynx, Bozeman, MT). A plastic core body, created by a 3D printer, contained multiple small holes with a dimeter of 0.7 mm distributed in space corresponding with the XY-coordinates of the targeted cortical areas, which served as a template to determine the locations of electrodes. An electrode assembly was created by setting metal tubes and electrodes into these holes so that the tips of the electrodes corresponded with the depth of individual brain regions. The other open ends of the electrodes were connected to metal holes of an EIB mounted on the top of a core body. An EIB consisted of 30 LFP channels and 2 ground channels, and all electrical signals from these channels were transferred to an Omentins connector. For brain LFP recording, a nichrome wire (A-M Systems, WA) or a tetrode that was constructed by bundling together four 17-μm polyimide-coated platinum–iridium (90/10%) wires (California Fine Wire, CA) and plated with platinum used to adjust the electrode impedances to 150–300 kΩ were used. The size of the electrode assembly was width 20 mm, length 20 mm, height 41 mm, and weight 2.2 g. The open edges of all electrodes were soldered to the corresponding channels on the EIB.

### Surgery

Standard surgical procedures were similar to those described previously (Konno et al., 2019; Sasaki et al., 2017; Shikano et al., 2018). Animals were anesthetized with 1–2% of isoflurane gas in air. The animal was then fixed in a stereotaxic instrument with two ear bars and a nose clamp. First, two craniotomies were made; one covering the coordinates for the anterior cingulate cortex (ACC; 1.2 mm anterior and 0.2 mm lateral to the bregma), primary motor cortex (M1: 1.2 mm anterior and 1.6 mm lateral to the bregma), and the striatum (STR: 1.2 mm anterior and 1.6 mm lateral to the bregma), and the other covering the coordinates for primary somatosensory cortex (S1; 1.9 mm posterior and 3.0 mm lateral to the bregma), the hippocampus (HPC; 1.9 mm posterior and 1.4 mm lateral to the bregma), and the medial parietal association cortex (MtPA; 1.9 mm posterior and 0.5 mm lateral to the bregma). The electrode array was directly implanted into the cortical tissue in the right hemisphere with electrodes inserted 0.5 mm into S1 and MtPA, 1.0 mm into ACC and M1, 1.4 mm into HPC, and 2.3 mm into STR. An electromyogram (EMG) electrode was implanted into the dorsal neck area. For the cerebellum, stainless steel screws were implanted on the skull attached to the brain surface to serve as ground/reference electrodes. Finally, all of the wires and the electrode array were secured to the skull using dental cement. After completing all surgical procedures, the anesthesia was terminated and the animals were spontaneously allowed to awake from the anesthesia. Following surgery, each animal was housed in a standard environment or an enriched environment with free access to water and food, with daily observation.

### In vivo electrophysiology

Before starting electrophysiological recording, the EIB on the animal’s head was connected to a digital headstage Cereplex M (Blackrock Microsystems, Salt Lake City, UT, U.S.A.), and the digitized signals were transferred to a data acquisition system Cereplex Direct (Blackrock Microsystems, UT). Local field potential (LFP) recordings commenced at a sampling rate of 2 kHz for 2.5–3.5 hours. Recordings were performed in an open-field box (25 cm wide × 18 cm deep × 13 cm high) located in a dark soundproofed room. For all recordings, electrophysiological signals were amplified and digitized at 2 kHz, and filtered between 0.1 Hz and 500 Hz.

### Histology of brain tissue

After the recordings, the mice were perfused intracardially with cold 4% paraformaldehyde (PFA) in 25 mM phosphate-buffered saline (PBS) and decapitated. The electrodes were carefully removed from the brain 6–8 hours after the perfusion. The brains were placed in 30% sucrose until equilibrated and coronally sectioned at a thickness of 50 mm, and the slices were stained with cresyl violet.

### Definition of awake/sleep states

All analyses were performed using custom-made MATLAB 2020a (MathWorks, MA, USA) and Python 3 routines. Awake/sleep states were defined from electromyography (EMG) signals. The root mean square (RMS) of the EMG signals was computed with a bin size of 1 s. The threshold to define awake/sleep states was set as the mean + 1.5 × standard deviations (SDs) for all the EMG RMS, and sleep states were defined as the periods when the RMS was below the threshold for more than 3 min. Awake states were defined as the other periods. Next, we classified sleep states into rapid eye movement (REM) sleep states and slow-wave sleep (SWS) states using EMG RMS and LFP power. Wavelet analysis was applied to LFP signals.

### LFP power and correlation analysis

LFP signals were downsampled to 400 Hz, and LFP spectral power in six frequency bands (δ: 1–4 Hz, θ: 4–8 Hz, α: 8–13 Hz, β: 13–30 Hz, low-γ: 30–50 Hz, and high-γ: 50–100 Hz) was computed by Morlet wavelet analysis. Each LFP power was normalized by the average during all sleep states in each frequency band. LFP signals from 2.5–3.5 hours were divided into bins of 10 s, and bins were categorized as awake bins when the bin contained an EMG RMS exceeding the threshold for more than 1 s. The other bins were categorized as sleep bins. In each bin, correlations of LFP power changes in each frequency band were computed from the 15 pairs of brain regions to obtain a 15-dimensional vector. When specified, vectors in all five frequency bands were combined to obtain a 90-dimensional vector.

### Machine learning algorithms

Uniform manifold approximation and projection (UMAP) was used for nonlinear dimensionality reduction ^35^. With UMAP, 15- or 90-dimensional vectors were reduced to two dimensions with the following parameters: *n_neighbors* = 3, *min_dist* = 0.1, *n_components* = 2, and *metric* = ‘euclidean’. Then, a robust continuous clustering (RCC) algorithm, an unsupervised clustering method, was applied to the plots from UMAP (Python implemented with the following parameters: *clustering_threshold* = 100, *k* = 100, and *measure* = ‘euclidean’) (Fig. 3) ^36^. If more than half of the bins in a cluster defined by the RCC were awake or sleep bins, the cluster was classified as an awake cluster or a sleep cluster, respectively. To quantify the degree of the separation between awake/sleep bins in awake/sleep clusters, in each animal, F1 scores for awake and sleep bins (termed F1_awake_ and F1_sleep_, respectively) were computed as the harmonic mean of precision and recall, where precision was the ratio of the number of awake/sleep bins included in awake/sleep clusters to the number of all bins included in the awake/sleep cluster, and recall was the ratio of the number of awake/sleep bins included in awake/sleep clusters to the number of all awake/sleep bins. In each animal, an F1 score was computed as (F1_awake_ + F1_sleep_)/2. To assess the significance of an F1 score, shuffled datasets were constructed for each original UMAP plot by randomly shuffling bin types (awake/sleep) assigned to the plots. An F1 score for the original data was considered to be significant when it was higher than all F1 scores computed from 1000 shuffled datasets.

### Gene expression analysis

For gene expression analysis, tissue samples were obtained from the dorsal hippocampus. Total RNA was extracted from the tissue using the RNeasy Plus Mini Kit (Qiagen Inc., Valencia, CA, USA). Microarray studies were performed using Affymetrix GeneChip Mouse Clariom S arrays (Thermo Fisher Scientific, Hampton, NH, USA). Gene expression analysis was performed using iDEP (integrated Differential Expression and Pathway analysis; Steven, Runan, 2018, BMC Bioinformatics).

### Statistics

All data are presented as the mean ± standard error of the mean (SEM), unless otherwise specified, and were analyzed using Python and MATLAB. Comparisons of two-sample data were analyzed by Student’s *t*-test and paired *t*-test and Mann Whitney U test. Multiple group comparisons of mean firing properties were performed by post hoc Bonferroni corrections. The null hypothesis was rejected at the *P* < 0.05 level.

### Data Availability

The datasets collected and analyzed are available from the corresponding author on request.

## Acknowledgement

This work was supported by KAKENHI (19H04897; 20H03545) from the Japan Society for the Promotion of Science (JSPS), a grant (JPMJCR21P1) from the JST CREST, and a grant (1041630; JP21zf0127004) from the Japan Agency for Medical Research and Development (AMED) to T. Sasaki; funds from the JST Exploratory Research for Advanced Technology (JPMJER1801), and Institute for AI and Beyond of the University of Tokyo to Y. Ikegaya.

## Author contributions

D.K. and T.S. designed the study. D.K acquired the electrophysiological data, performed the analyses, and prepared the figures. D.K. and T.S. wrote the main manuscript text. Y.I. supervised the project, and all the authors reviewed the main manuscript text.

---

**Figure S1.**
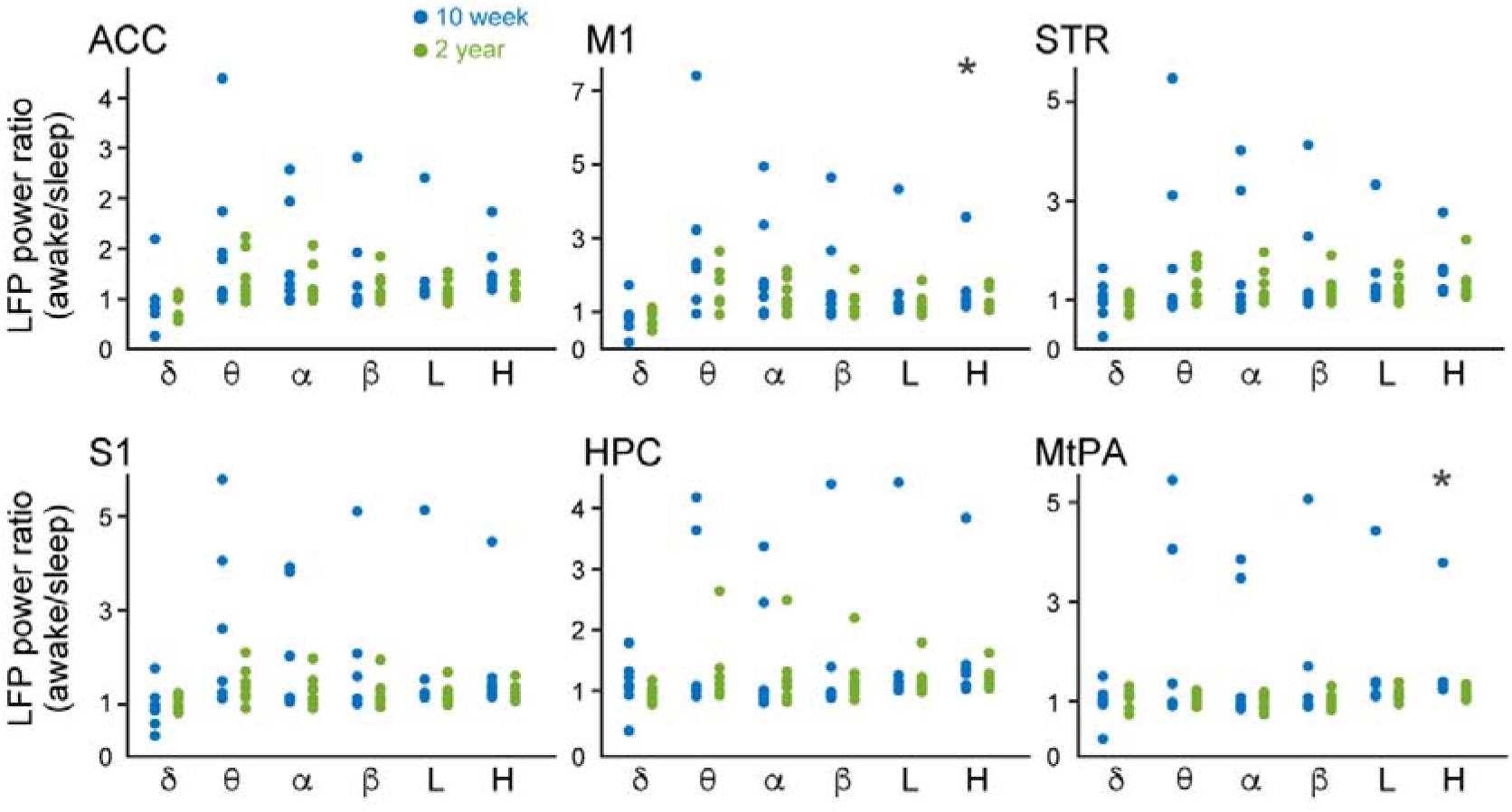
LFP power in single frequency bands in single brain regions in awake and sleep states. LFP power in individual frequency bands in individual brain regions in awake states was normalized by the average in sleep states (*n* = 7 and 8 mice). \**P* < 0.05, Student’s *t*-test versus 1 in each group.

**Figure S2.**
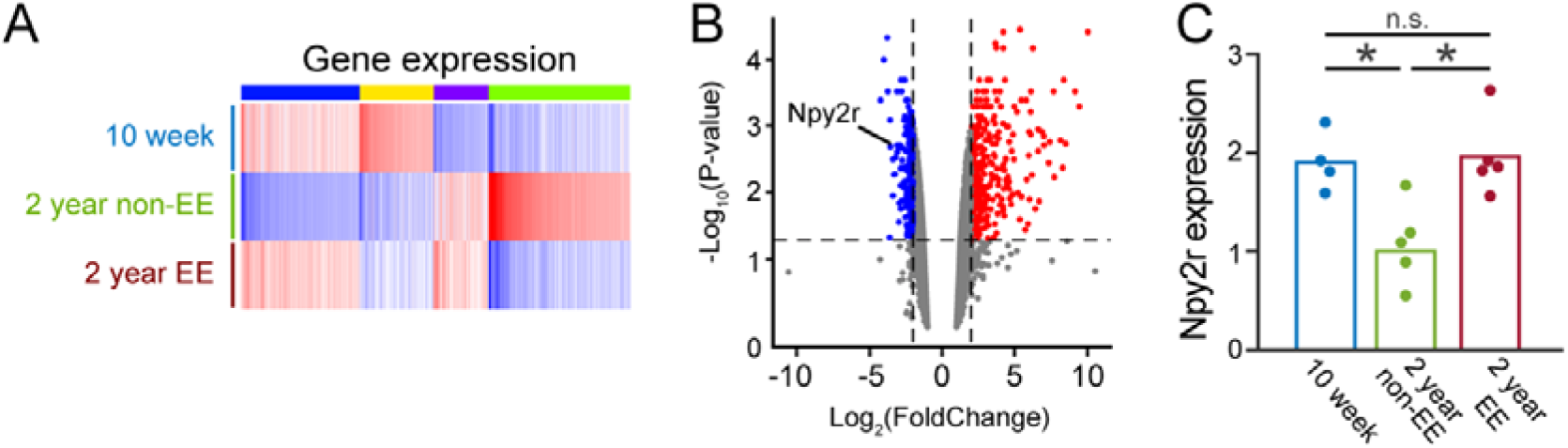
Gene expression in 2-year EE mice. **(A)** Heatmaps of representative gene expression patterns in the dorsal hippocampus of 10-week, 2-year non-EE, and 2-year EE mouse groups. Of the 21200 genes, the top 2000 genes with large expression variations between groups are shown. The top color bar shows clusters defined by the *K*-means algorithm. The gene ontology of the genes in each cluster is shown in Figure S3. **(B)** Volcano plot showing the fold change and *P* value of individual genes in 2-year non-EE mice compared with 10-week mice. The vertical dotted lines indicate 2-fold upregulation and downregulation, and the horizontal dotted line represents the cutoff significance level (*P*□=□0.05). The red and blue dots indicate significantly upregulated and downregulated genes, respectively, in aged mice. **(C)** qPCR analysis validating the expression levels of Npy2r, a gene found to be differentially expressed in the bulk RNA-seq analysis across the mouse groups (*n* = 4, 5, and 5 mice). Each dot represents an individual mouse. \**P* < 0.05, Student’s *t*-test.

**Figure S3.**
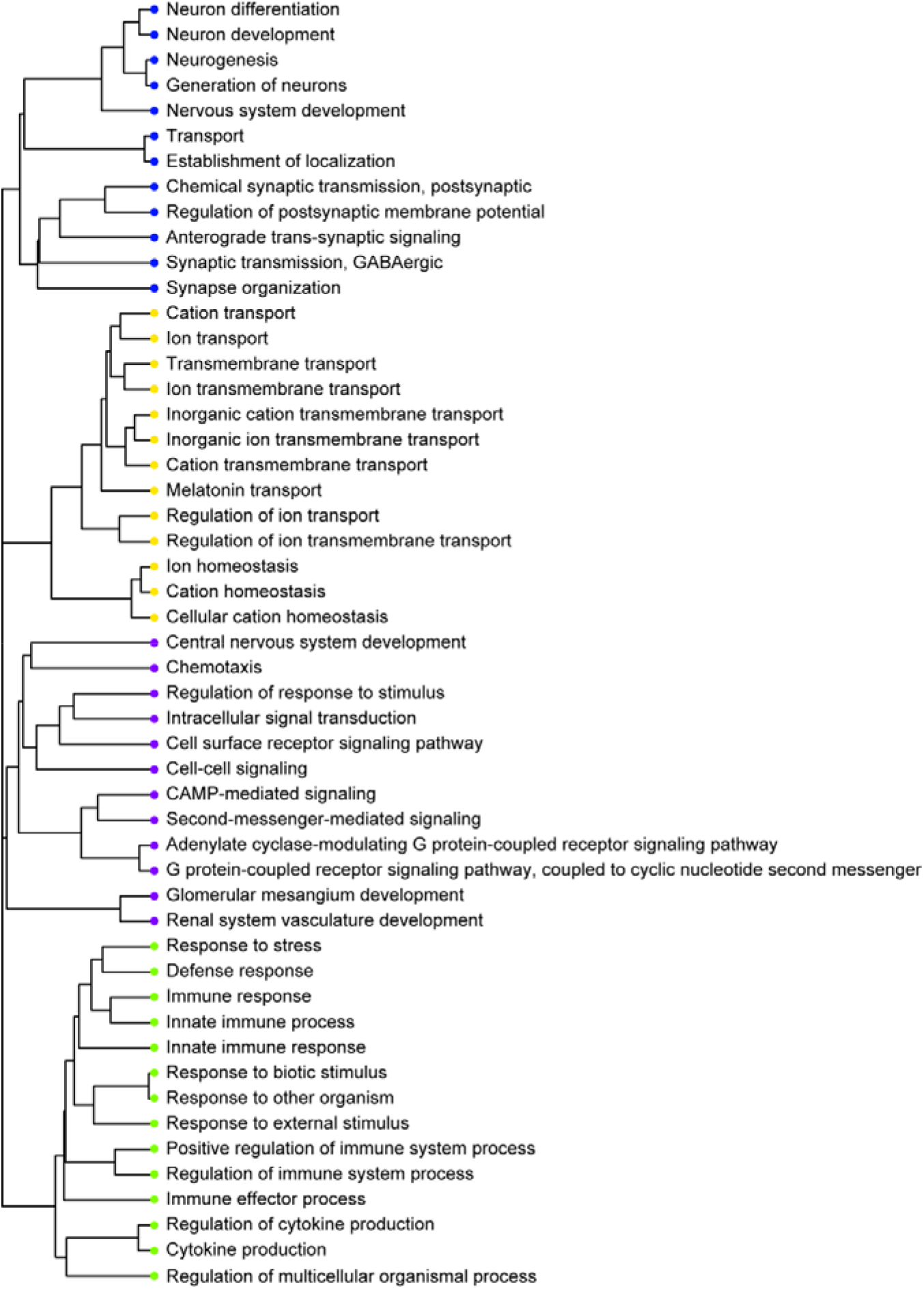
Gene Ontology (GO) enrichment analysis for each cluster of the heatmap shown in Figure S2A. The cluster with increased expression in the 2-year non-EE group (green) contains GO terms related to stress and immune responses, while the cluster with increased expression in the 10-week and 2-year EE groups (blue) contains GO terms related to neurogenesis and synaptic transmission.

